# Determining the inherent reaction-diffusion properties of actin-binding proteins in cells by incorporating genetic engineering to FRAP-based framework

**DOI:** 10.1101/2020.09.21.305615

**Authors:** Takumi Saito, Daiki Matsunaga, Tsubasa S. Matsui, Kentaro Noi, Shinji Deguchi

## Abstract

Proteins in cells undergo repeated association to other molecules, thereby reducing the apparent extent of their intracellular diffusion. While much effort has been made to analytically decouple these combined effects of pure diffusion and chemical reaction, it is difficult to attribute the measured quantities to the nature of specific domains of the probed proteins particularly if, as is often the case, the protein has multiple domains to independently interact with the same types but different molecules. Motivated by the common goal in cell signaling research aimed at identifying the protein domains responsible for particular intermolecular interactions, here we describe a new approach to determining the domain-level reaction and pure diffusion properties. To validate this methodology, we apply it to transgelin-2, an actin-binding protein whose intracellular dynamics remains elusive. We develop a fluorescence recovery after photobleaching (FRAP)-based framework, in which comprehensive combinations of domain-deletion mutants are created with genetic engineering, and the difference among the mutants in FRAP response is analyzed. We demonstrate that transgelin-2 in cells interacts with F-actin via two separate domains, and the chemical equilibrium constant of the interaction is determined at the individual domain levels. Its pure diffusion properties independent of the association to F-actin is also obtained. This approach requires some effort to construct the mutants, but instead enables in situ domain-level determination of the physicochemical properties, which will be useful, as specifically shown here for transgelin-2, in addressing the signaling mechanism of cellular proteins.

## Introduction

Proteins undergo repeated association and dissociation to their partner proteins in cells, a universal event called “turnover” (Smith et al. 2013). Because of this physicochemical interaction among proteins, their transport within cells is dominated not only by the Brownian motion-based pure diffusion but also by the molecular turnover. The pure diffusion of particles is characterized by the diffusion coefficient *D*= ⟨*r*^2^⟩/4*t* where ⟨*r*^2^⟩ is the mean square displacement (MSD) over time *t*. The extent of the association and dissociation of proteins is typically characterized by the association rate *k*_on_ and dissociation rate *k*_off_. Pure diffusion corresponds to a case where *k*_on_ = 0, but in reality the intracellular diffusion of proteins is more or less affected by the binding to other molecules, and thus it is not straightforward to decouple the mixed effects within cells into each of the inherent reaction and diffusion properties specific to the proteins.

One possible way to circumvent the above issue might be evaluating the association and dissociation rates in vitro by using, for example, the stopped-flow method (e.g., Goldmann and Isenberg 1993; Hundt et al. 2016). However, the reaction rates measured in extracellular environments are not always consistent with those in cells particularly because there are in general kinetic competitions among multiple proteins, while it is hard to accurately mimic the intracellular milieu in in vitro experiments. Besides, a specific protein of interest may form a complex with other molecules, which can significantly alter the kinetic properties (Nakorchevsky et al. 2010; Matsui et al. 2018).

To instead directly characterize the kinetics of proteins within cells, a microscopic technique called fluorescence recovery after photobleaching (FRAP) is often used, in which the temporal evolution of fluorescence-labeled proteins is analyzed during the bleaching and recovery (Blonk et al. 1993; Daddysman and Fecko 2013). The fluorescence recovery curve is often fitted by single/double exponential functions, and then the recovery rate is determined from the inverse of the characteristic time constant (Campbell and Knight 2007; Sakurai-Yageta et al. 2015; Dukic et al. 2017). Otherwise, more elaborate physical models have been employed to separately evaluate the reaction-diffusion properties of proteins in cells (Sprague et al. 2006; Mueller et al. 2008). However, the interpretation of the model output could be complicated if the probed protein has, as is often the case, multiple domains that are independently able to interact with the same types but different molecules. Particularly in the field of cell signaling research, it is essential to identify which domains within the individual proteins are responsible for specific intermolecular interactions. Thus, it is hard in conventional approaches to attribute the measured mixed kinetics to the nature of specific domains of the probed proteins. Another drawback of previous approaches is that, to estimate reaction-free pure diffusion properties within cells, the biologically inert green fluorescent protein (GFP) is often employed as a substitute of a target protein, but the molecular weight that affects the diffusion may not necessarily be close to that of the actual one.

Here we present an alternative approach to determining the pure diffusion properties and intracellular kinetics of actin-binding proteins (ABPs). To validate the methodology, we apply it to transgelin-2 (Leung et al. 2011; Na et al. 2015; Kim et al. 2018) – a 22-kDa ABP and also known as smooth muscle protein 22β or SM22β – whose intracellular dynamics remains elusive particularly due to the presence of two potential actin-interacting domains. Genetic engineering is incorporated to construct the mutants of transgelin-2, and the difference among the mutants in the spatiotemporal response in FRAP experiments is analyzed. We then extract the pure diffusion coefficient intrinsic to transgelin-2 as well as its dissociation constant from the actin filaments, in which the latter is determined for each of the actin-interacting domains. The approach described here will be useful in the domain-level characterization of the physicochemical properties of cellular proteins.

## Materials and methods

### Cells, plasmids, and antibodies

Rat aortic smooth muscle cell lines (A7r5, ATCC) were cultured with low-glucose (1.0 g/L) Dulbecco’s Modified Eagle Medium (Wako) containing 10% (v/v) heat-inactivated fetal bovine serum (SAFC Biosciences) and 1% penicillin-streptomycin (Wako) in a 5% CO_2_ incubator at 37°C. Expression plasmids encoding mClover2-tagged transgelin-2 (TAGLN2) and mRuby2-tagged Lifeact were constructed by inserting the PCR-amplified cDNAs (human TAGLN2, pFN21ASDA0120, Kazusa DNA Research Institute; Lifeact, Addgene plasmid # 54688; a gift from Michael Davidson) into the mClover2-C1 vector (Addgene plasmid #54577, a gift from Michael Davidson) and the mRuby2-N1 vector (Addgene plasmid #54614, a gift from Michael Davidson), respectively. The following domain-deletion mutants of TAGLN2 were constructed by inverse PCR using a KOD Plus Mutagenesis Kit (Toyobo) (Table S1 for the sequences of the primers used): ΔCH (Δ25-136), ΔAB (Δ153-160), ΔCR (Δ174-197), ΔCHΔCR, ΔABΔCR, ΔABΔCH, and ΔABΔCHΔCR. An expression plasmid encoding mClover2-beta-actin was constructed by inserting a human beta-actin gene, which was digested with XhoI and BamHI restriction enzymes from the EYFP-actin vector (#6902-1, Clontech) into the mClover2-C1 vector. These plasmids were transfected to cells 24 h after the seeding using Lipofectamine LTX and plus Reagent (Thermo Fischer Scientific) according to the manufacturer’s instructions. Rabbit polyclonal anti-TAGLN2 (ab121146, Abcam), mouse monoclonal anti-GFP (012-22541, Wako), and anti-beta-actin (G043, ABM) were used for detecting endogenous TAGLN2 proteins, mClover2 tagged-mutants of TAGLN2, and beta-actin proteins, respectively.

### Western blotting

Cells transfected with the plasmids for 24 h were lysed by the SDS sample buffer. Lysates were centrifuged at 14,000 rpm for 15 min at 4 □, and the supernatants were collected. Protein concentration was quantified by BCA Protein Assay Kit (Thermo Fischer Scientific). The proteins were fractionated by 12% gradient SDS-PAGE, transferred onto PVDF membranes (Millipore), blocked with 5% (w/v) BSA for 1 h, incubated with the primary antibodies overnight, blocked again with 4% skim milk for 1 h, and incubated with the HRP-conjugated secondary antibodies (Bio-Rad; #1705047 for β-actin and mClover2, #1705046 for TAGLN2) for 1 h and with the enhanced chemiluminescence reagent (Immobilon Western, Millipore) for 1 h. Bands were detected by Immobilon Western (Millipore). Chemiluminescence images were taken by CCD-based imaging system (ChemiDoc XRS+, Bio-Rad).

### FRAP experiments

Cells were cultured on a glass-bottom dish and transfected with the plasmids for 24 h for FRAP experiments. The fluorescence intensity profiles were obtained by using the FV1000 confocal laser scanning microscope (Olympus) with a 60X oil immersion objective lens (NA = 1.42). A square region containing a single SF was bleached for 1 s by using a 405/440-nm wavelength laser. Meanwhile, the line-scanning was conducted along the longitudinal length every 1.62 ms for totally ∼10 s (to acquire in total 6,000 frames) using a 488-nm wavelength laser. Setting *t* = 0 s to be the time of completion of the photobleaching, ventral stress fibers (SFs) were thus scanned during pre-bleach (−2 ≤ *t* < −1 s), photobleaching (−1 ≤ *t* < 0 s), and recovery (*t* ≥ 0 s). 2D-scanning was then followed to capture the entire bleached regions every 0.5 s for ∼30 s (to acquire in total 60 frames). Images were analyzed using ImageJ software (NIH) and MATLAB (MathWorks). The time evolution of the fluorescence intensity spatially averaged over the bleached length *l*_X_ (see also Fig. 4A), normalized by the original intensity averaged temporally over the period of pre-bleach and spatially over the same length *l*_x_, is fitted by the least-square method to a single exponential equation

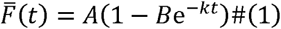

where *k* (=1/*τ*) represents the recovery rate or the inverse of the time constant *τ*, and *A* and *B* are fitting parameters; specifically, *A* represents the mobile fraction.

**Fig. 1.**
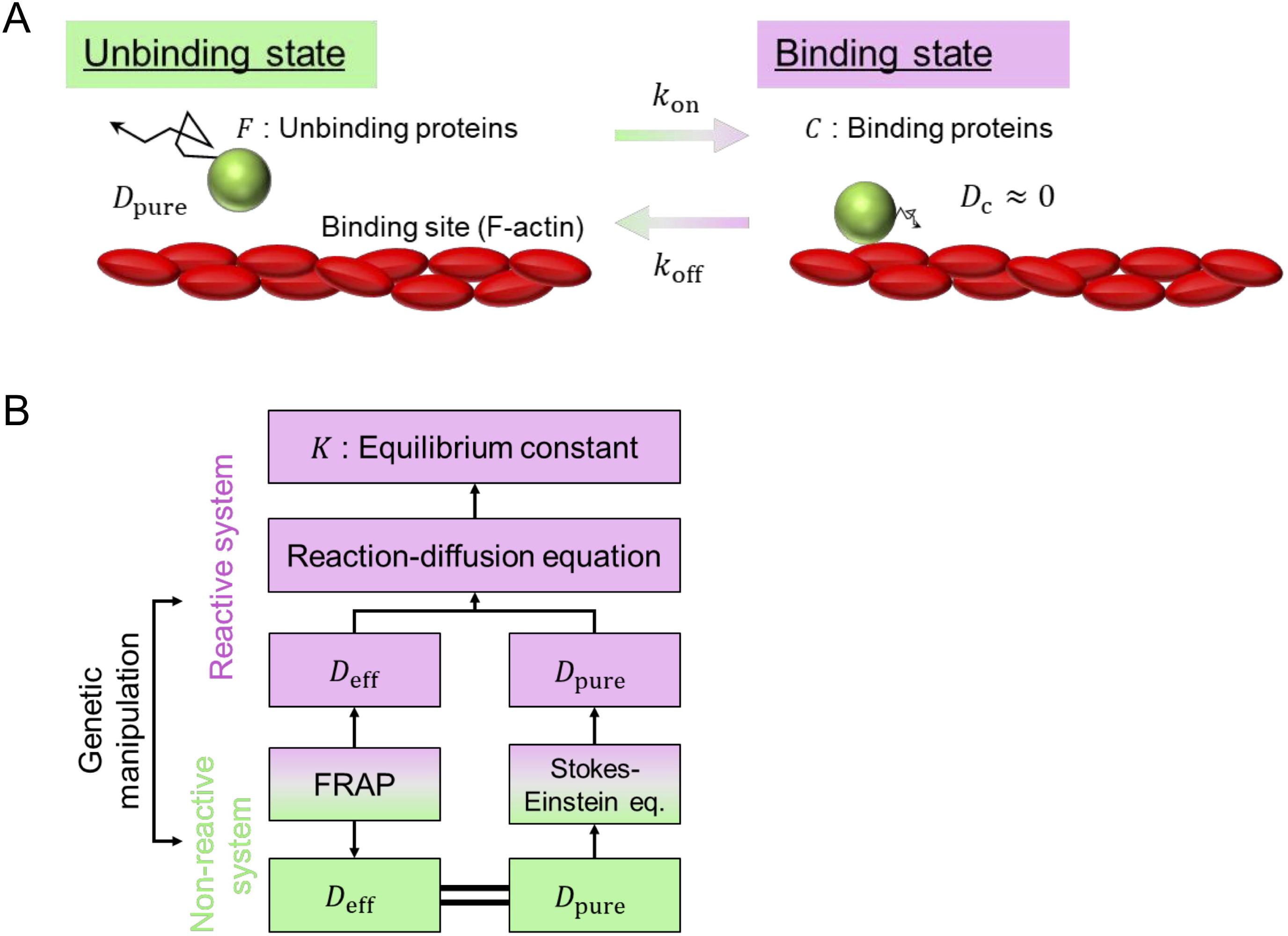
Scheme of the research design. (A) Variables and parameters related to the binding and unbinding states of ABPs. *F* and *C* represent the fluorescence intensity of each condition. (B) Strategy to determine the reaction-diffusion properties of ABPs. Genetic manipulation is employed to produce reactive and non-reactive systems. *D*_eff_, which is obtained by FRAP experiments, is identical to *D*_pure_ in the non-reactive system and is in turn converted to *D*_pure_ at the reactive system with modification according to the Stokes-Einstein equation. The equilibrium dissociation constant in the reactive system *K* is then determined through its relationship with *D*_eff_ and *D*_pure_ to be in line with the reaction-diffusion equation. See the text for further details.

**Fig. 2.**
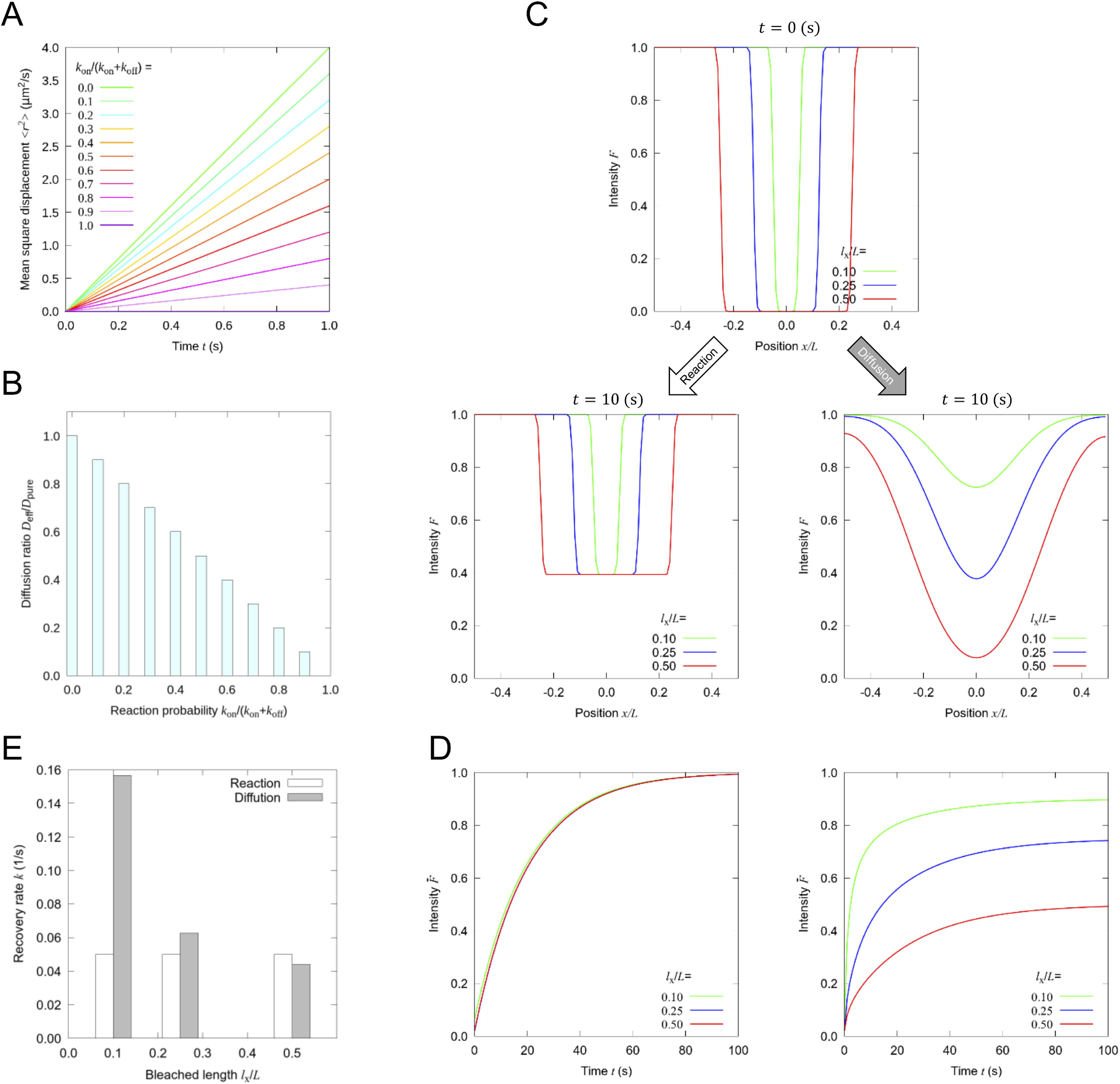
Numerical analyses to support the research design. (A) MSD of randomly walking particles vs. elapsed time as a function of the given reaction probability parameter *k*_on_/(*k*_on_ + *k*_off_), in which 0 and 1 result in pure diffusion and immobilization, respectively. (B) *D*_eff_/*D*_pure_ is linearly decreased with the reaction probability parameter *k*_on_/(*k*_on_ + *k*_off_). (C) The recovery process of FRAP is simulated by two distinct models, i.e., a completely reaction-driven (Eq. (11)) or diffusion-driven model (Eq. (10)), with a parameter of the bleached length *l*_x_ normalized by the region of interest *L* (or the experimentally scanned region). (D) Time course of the fluorescence recovery with the completely reaction-driven (left) or diffusion-driven (right) model. (E) Recovery rate *k* as a function of the normalized bleached length *l*_x_/*L* obtained by fitting Eq. (1) to the data in D with the least-square method.

### Model description

We consider general ABPs (Fig. 1A), for which we later assume a specific protein – transgelin-2 encoded by the TAGLN2 gene. The reaction-diffusion equation of the system is described by

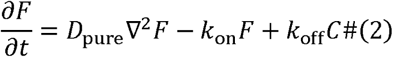

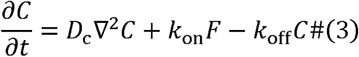

where *F* and *C* represent the concentrations of an ABP at the unbinding (free) and binding (complex) state, respectively; *D*_pure_ and *D*_C_ represent the pure diffusion coefficients at the unbinding and binding state, respectively; *k*_on_ and *k*_off_ represent the association and dissociation rates, respectively; ∇^2^ represents the Laplace operator. We now introduce two assumptions in accordance with previous studies dealing with similar systems (Sprague et al. 2004a). First, the binding site of the actin filaments is considered immobile, and thus the diffusion is ignored upon the binding (i.e., *D*_c_ ≈ 0). This assumption is reasonable for the present case as the turnover of the actin filaments in SFs is much slower than that of transgelin-2 as demonstrated in the results. Second, the reaction is considered to take place under equilibrium, which is reasonable as the total amounts of transgelin-2 and actin will remain constant over the brief time course of the FRAP experiments, and in addition photobleaching is supposed to affect only the fluorescence intensity but not the reaction rate. Consequently, during the binding state, the system is virtually at a steady state, and hence Eq. (3) is reduced to

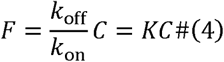

where *K* represents the equilibrium dissociation constant defined as *K* = *k*_off_ / *k*_on_. Thus, Eq. (2) is now reduced to a form of the diffusion equation

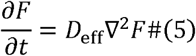

where

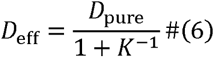

characterizes the slowed diffusion of the ABP due to its binding to the actin filaments in SFs, which has been referred to as “effective” diffusion coefficient (Crank 1975; Sprague et al. 2004a; Ait-Haddou et al. 2010). The analytical solution of Eq. (5) is described on a rectangular bleached region by

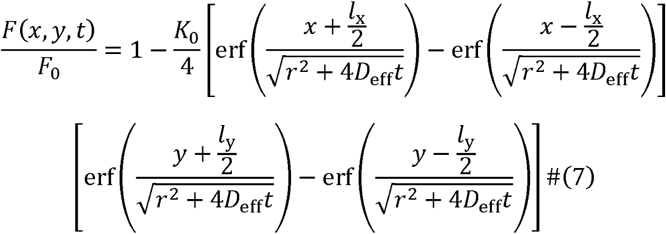

where *F*_0_, *K*_0_, *r, l*_x_, and *F*_y_, and erf represent the intensity in the pre-bleach state, photobleaching parameter, laser resolution, width and height of the bleached rectangular area, and the error function (i.e., 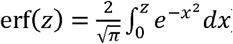), respectively (Deschout et al. 2010).

### Determination of the reaction-diffusion parameters

FRAP experiments that provide the effective diffusion coefficient *D*_eff_ are per se not sufficient for determining the pure diffusion coefficient *D*_pure_. To overcome this limitation, deletion mutants of the protein with (reactive system) or without (non-reactive system) actin-binding domains are constructed by genetic manipulation (Fig. 1B). The effective diffusion coefficient obtained with the FRAP on the non-reactive system is regarded as the pure diffusion coefficient of the system, i.e., *D*_eff_ = *D*_pure_. The actual pure diffusion coefficient in the reactive system is estimated from the pure diffusion coefficient in the non-reactive system of ΔABΔCH by compensating for the effect of the molecular weight (*M*) according to the Stokes-Einstein equation, 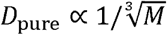 for spherical particles with constant density (Sprague et al. 2004b). Together with this pure diffusion coefficient and the effective diffusion coefficient obtained in the reactive system, the equilibrium dissociation constant in the reactive system is now estimated following Eq. (6). The spatiotemporal intensity data in the FRAP experiments were then applied to Eq. (7) to determine the values of the reaction-diffusion parameters by the maximum likelihood method (Deschout et al. 2010).

### Numerical analysis: Brownian dynamics simulation of reaction-diffusion systems

2D numerical analyses were performed to describe how the diffusion of particles is contaminated by the chemical interaction with the surroundings characterized by *k*_on_ and *k*_off_. Stochastic events of either binding or unbinding are determined at each time step by a number *P* between 0 and 1 from a random number generator, at which particles with *P* ≥ *k*_on_/(*k*_on_ + *k*_off_) are bound to hypothetical immobile sites, whereas those with *P* < *k*_on_/(*k*_on_ + *k*_off_) are allowed to move to follow the random Gaussian process with the displacement variance *σ*^2^ = 2*D*_pure_Δ*t* where Δ*t* represents the time interval (Qian et al. 1991; Nitsche and Hinch 1995). The resulting MSD and the effective diffusion coefficient at time *t* were computed by

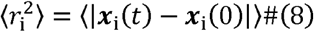

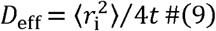

where *x*_*i*_ is the position of the *i*-th particle (Qian et al. 1991; Smelser et al. 2015; Matsunaga et al. 2020).

### Numerical analysis: continuum simulation of two extreme cases of the FRAP response

1D numerical analyses were performed to highlight the difference in the response in FRAP experiments between pure diffusion-based recovery and turnover-based one. Temporal change in the concentration of fluorophores *F* is described in the pure diffusion-driven case by

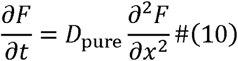

or in the turnover-driven case by

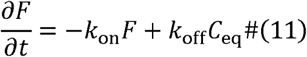

where *C*_eq_ is the equilibrium concentration of a reaction partner *C* under an assumption that *C* is virtually unchanged in concentration over time. The initial configuration of the fluorophores in the bleached region is determined by substituting *t* = 0 to the 1D equivalent of Eq. (7) as

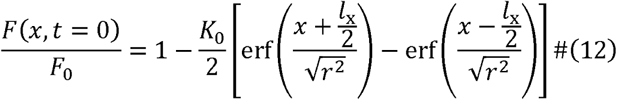

and subsequent temporal evolution was analyzed by Eq. (10) or by Eq. (11), with using a forward difference in time and a central difference in space (Larkin 1964).

### Fluorescence correlation spectroscopy

Fluorescence correlation spectroscopy (FCS) was performed using the FV1000 confocal laser scanning microscope with a 60X oil immersion objective lens (NA = 1.42) at the outside or inside of the cells cultured on a glass-bottom dish and transfected with the plasmid of the non-reactive ΔABΔCH of TAGLN2 for 24 h. The diffusion coefficient *D* and the average particle number 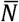 were determined by the least-square method from the autocorrelation function *G* of the fluorescence intensity *I*(*t*) as a function of the lag time *τ* described as

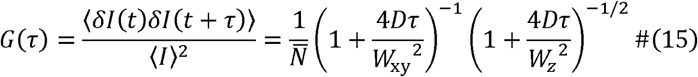

where *δ I*(*t*) (= *I*(*t*) − ⟨ *I* ⟩) represents the fluctuation of the intensity detected every 2 ns, and *W*_xy_ and *W*_z_ represent confocal volume parameters (0.17 μm and 2.8 μm, respectively) determined from the current setup of the numerical aperture, wavelength, and pin-hole size (Krichevsky and Bonnet 2002).

### Statistical analysis

Unless otherwise stated, data are expressed as the mean ± standard deviation of more than 5 independent experiments. Differences were calculated based on the unpaired Student’s *t*-test, with a significance level of p < 0.01 (*) or p < 0.05 (**).

## Results

### Numerical simulations validate the research design

To illustrate the conceptual basis of the present study, we simulated 2D diffusion of particles undergoing stochastic interactions with hypothetical immobile sites. The MSD was analyzed by tracking the position of individual particles to show that the MSD is proportional to the elapsed time as expected (Fig. 2A; Table S2 for the parameters used). The slope of the MSD–time curve, corresponding to the diffusion coefficient, was confirmed to be smaller with larger binding probabilities. As such, the ratio of the diffusion coefficient at specific binding probability (i.e., effective diffusion) to that at *k*_on_ = 0 (i.e., pure diffusion), namely *D*_eff_/*D*_pure_, was linearly decreased upon the increased binding probability (Fig. 2B). Thus, the substantive diffusion is slowed in the presence of the chemical interaction, which we aim with the experiments below to decompose into the two independent factors, i.e., the inherent pure diffusion and chemical interaction.

We also numerically simulated the response in FRAP experiments to validate our approach to determining the above two factors. Specifically, our approach is essentially based firstly on the use of Eq. (7) and secondly on that of the genetic engineering (Fig. 1B). Regarding this first basis, we use below both the spatial and temporal information in Eq. (7) to interpret the FRAP data, while the single exponential approach of Eq. (1) – or more generally an approach omitting the spatial information – should be inadequate in the present aim. To numerically demonstrate this, the recovery process along a bleached length *l*_x_ driven by the pure diffusion according to Eq. (10) was compared to that driven completely by the chemical reaction according to Eq. (11) (Fig. 2C; Table S3 for the parameters used). The intensity profile dominated by the chemical reaction is spatially uniformly recovered while maintaining the initial form, whereas that dominated by the diffusion displays a normal distribution. The time-course change at the bleached region consequently exhibits an identical curve regardless of the bleached length *l*_x_ in the chemical reaction-dominant case, whereas that in the diffusion-dominant case does depend on *l*_x_ (Fig. 2D). Thus, except such a fully chemical reaction-dominant case, the recovery rate *k* (Kapitza et al. 1985; Chloë Bulinski et al. 2001; Sprague et al. 2004a) determined using Eq. (1) or the extent of pure diffusion is underestimated particularly with a large-sized *l*_x_ (Fig. 2E). This limitation on the dependence on the bleach size is circumvented by analyzing the spatial distribution in Eq. (7).

Regarding the above-described second basis of the necessity of the genetic engineering, the diffusion coefficient determined with Eq. (7) is only an “effective” one, to which the chemical reaction inevitably simultaneously contributes. To isolate the respective contributions, the reactive domain(s) in the protein is identified and deleted by genetic manipulation, and the resulting deletion mutant is tested for determining the now “pure” diffusion properties as the protein mutant is biologically inert and thus behaves only in a diffusive way (Fig. 1B). Finally, *K* that characterizes the chemical interaction in Eq. (4) is extracted from Eq. (6), in which the pure diffusion coefficient is modified by compensation of the decreased mass due to the domain deletion.

### Transgelin-2 interacts with F-actin in living cells via two actin-binding domains

Trangelin-2 – containing an N-terminal calponin homology domain (CH domain) between 25 to 136 amino acid residues, an actin-binding motif (AB domain) between 153 to 160 amino acid residues, and a C-terminal calponin-repeat domain (CR domain) between 174 to 197 amino acid residues – has been reported to be associated with F-actin via the CH and AB domains (Castresana and Saraste 1995; Solway and Fredberg 1997; Na et al. 2015). To modulate the association with F-actin, we constructed comprehensive combinations of deletion mutants of transgelin-2 tagged with mClover2 at the N-terminus, in which the CH, AB, and/or CR domains were deleted (Fig. 3A). We expected that the mutants with at least one of the CH and AB domains, in addition to the full-length wild type (FL), are able to be associated with F-actin (referred to as the reactive system), while those without both of the two domains are unable (the non-reactive system).

**Fig. 3.**
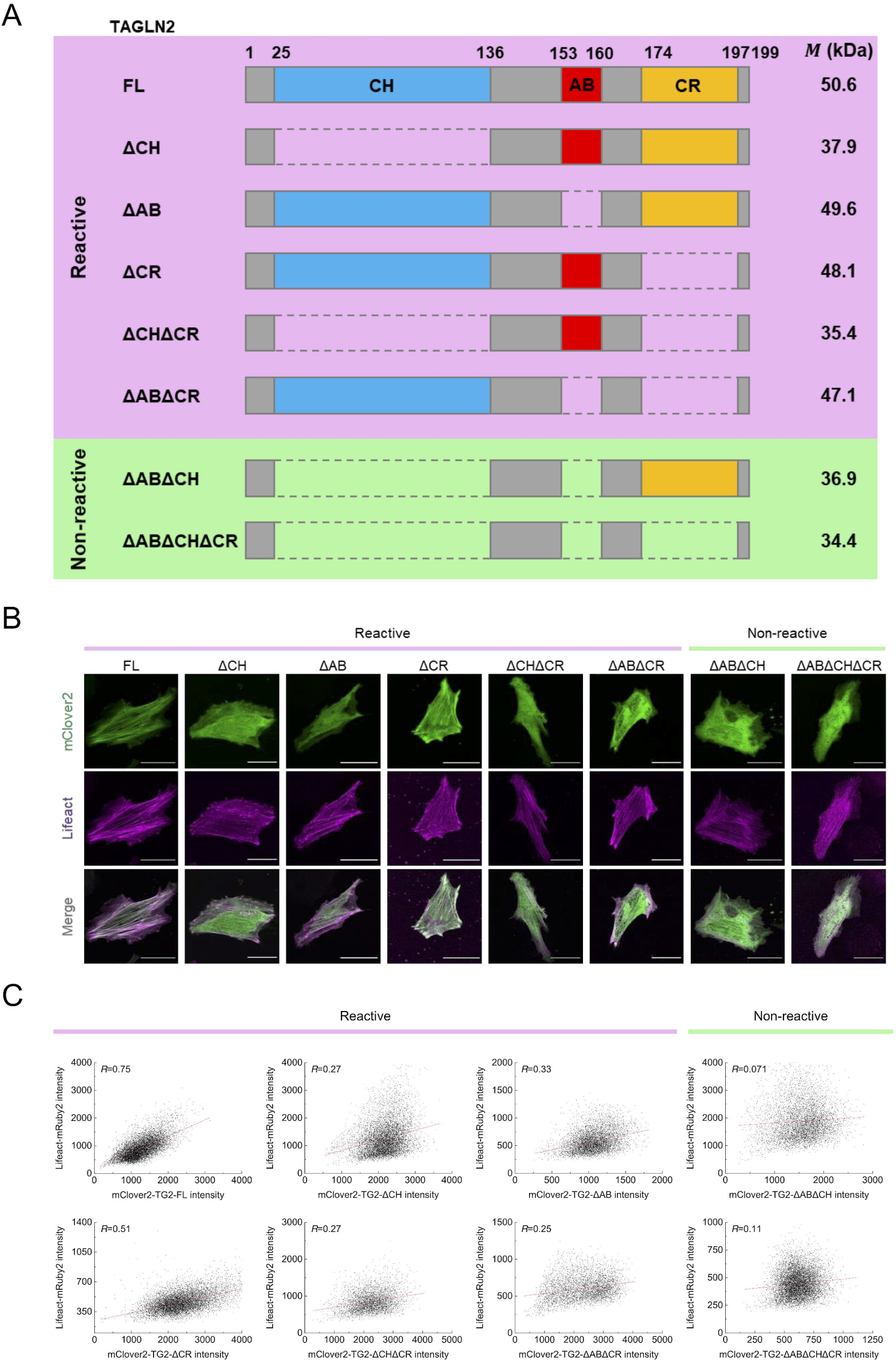
Transgelin-2 binds F-actin via the CH and AB domains. (A) Schematic diagram of the domain structure of transgelin-2 (encoded by the 7.4 □ kb TAGLN2 gene) and its mutants. mClover2 present at the N-terminus is omitted. (B) Confocal images of representative cells to analyze the intracellular colocalization between transgelin-2 or its mutant (mClover2) and F-actin (Lifeact). Scale, 20 μm. (C) The relationship of the fluorescence intensity between Lifeact (ordinate) and mClover2 (abscissa) measured at each pixel of the confocal images, with Pearson correlation coefficient *R* between the two fluorescent labels.

**Fig. 4.**
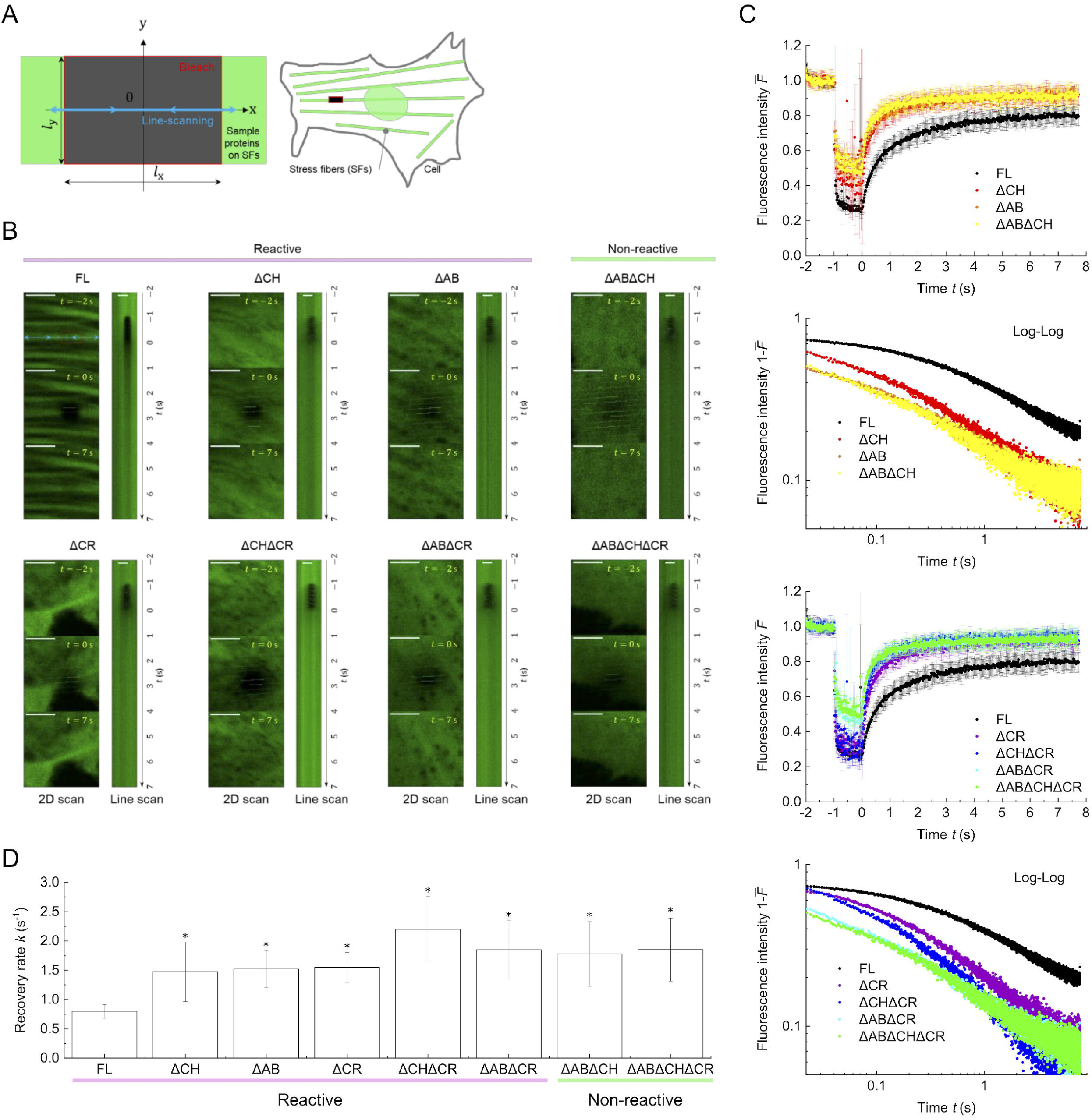
FRAP analysis suggests that the deletion of the domains of transgelin-2 destabilizes its association to F-actin. (A) A part of individual SFs is bleached in 2D, and the resulting fluorescence recovery is probed by the line-scanning along the middle of the bleached region at a high temporal resolution. (B) Confocal images of transgelin-2 and its mutants taken before (*t* = -2 s), just after (*t* = 0 s), and after (*t* = -7 s) the bleaching, with respective kymographs on the right. Scale, 1 μm. (C) Time course of 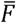 for different mutants is shown by two ways: a linear scale plot (upper) and a log-log plot (lower). (D) *k* for different mutants. Asterisks represent statistically significant differences compared to FL (*, p< 0.01).

To confirm the colocalization with F-actin, A7r5 cells transfected with one of the mClover2-labeled mutants and Lifeact-mRuby2 that visualizes F-actin were imaged and analyzed (Fig. 3B, 3C). FL and ΔCR clearly displayed a SF-like fibrous pattern similar to that of Lifeact-mRuby2, with a high level of Pearson correlation coefficient *R* between the two fluorescent labels of 0.75 and 0.51, respectively. With the deletion of either the CH or AB domain (i.e., ΔAB, ΔABΔCR, ΔCH, and ΔCHΔCR), the transgelin-2 mutants displayed a relatively diffusive but slightly fibrous pattern, with a moderate *R* level of 0.25–0.33. With the deletion of both ΔCH and ΔAB (i.e., ΔABΔCH and ΔABΔCHΔCR), the transgelin-2 mutants displayed a completely diffusive pattern with a low *R* level below 0.11, while Lifeact-mRuby2 or F-actin in SFs still exhibited the fibrous pattern. Thus, at least the AB or CH domain is indeed required for the association of transgelin-2 to F-actin, while the CR domain is dispensable.

### F-actin in SFs works as an immobile scaffold for transgelin-2

In the FRAP experiments using the transgelin-2 mutants, line-scanning was conducted along the center of the bleached region to detect the fluorescence recovery at a high temporal resolution (Fig. 4A). FL, as a control, exhibited a slow recovery and a large immobile fraction compared to the other mutants (Fig. 4B, 4C; Video S1), suggesting that the association to F-actin impedes the substantive diffusion of transgelin-2. The time evolution of the fluorescence was analyzed based on the single exponential function of Eq. (1), and the recovery rate *k* (Fig. 4D) and mobile fraction *A* (Fig. S1) were quantified. We found that *k* was significantly increased with any deletion mutant up to ∼1.5–2.0 s^-1^ on average compared to the control FL with an average *k* of ∼0.75 s^-1^ (Fig. 4D, Video S2–S8). ΔCR thus resulted in an increase in *k* compared to FL, although this mutant possesses both of the CH and AB domains used for the association to F-actin. This increase might be caused because the removal of the CR domain distorts the rest of the protein conformation including the critical CH/AB domains to reduce its affinity to F-actin, and/or decreases the molecular weight to increase the pure diffusion coefficient. Similar tendency was obtained for the mobile fraction (Fig. S1). Notably, no significant difference was observed among the deletion mutants (Fig. 4D, S1), but as described in the next section the current approach based on Eq. (1) may be inadequate to accurately capture the potential functional differences among them.

Nevertheless, in a separate FRAP experiment, the turnover of F-actin tagged with mClover2 was found to exhibit a recovery rate *k* of 0.0095 ± 0.0077 s^-1^, i.e., two orders of magnitude lower than that of transgelin-2 (Fig. 5A, 5B; Video S9). These results suggest that F-actin in SFs may be regarded as an immobile scaffold, and the assumption introduced to set *D*_c_ ≈ 0 in Eq. (3) is justified.

**Fig. 5.**
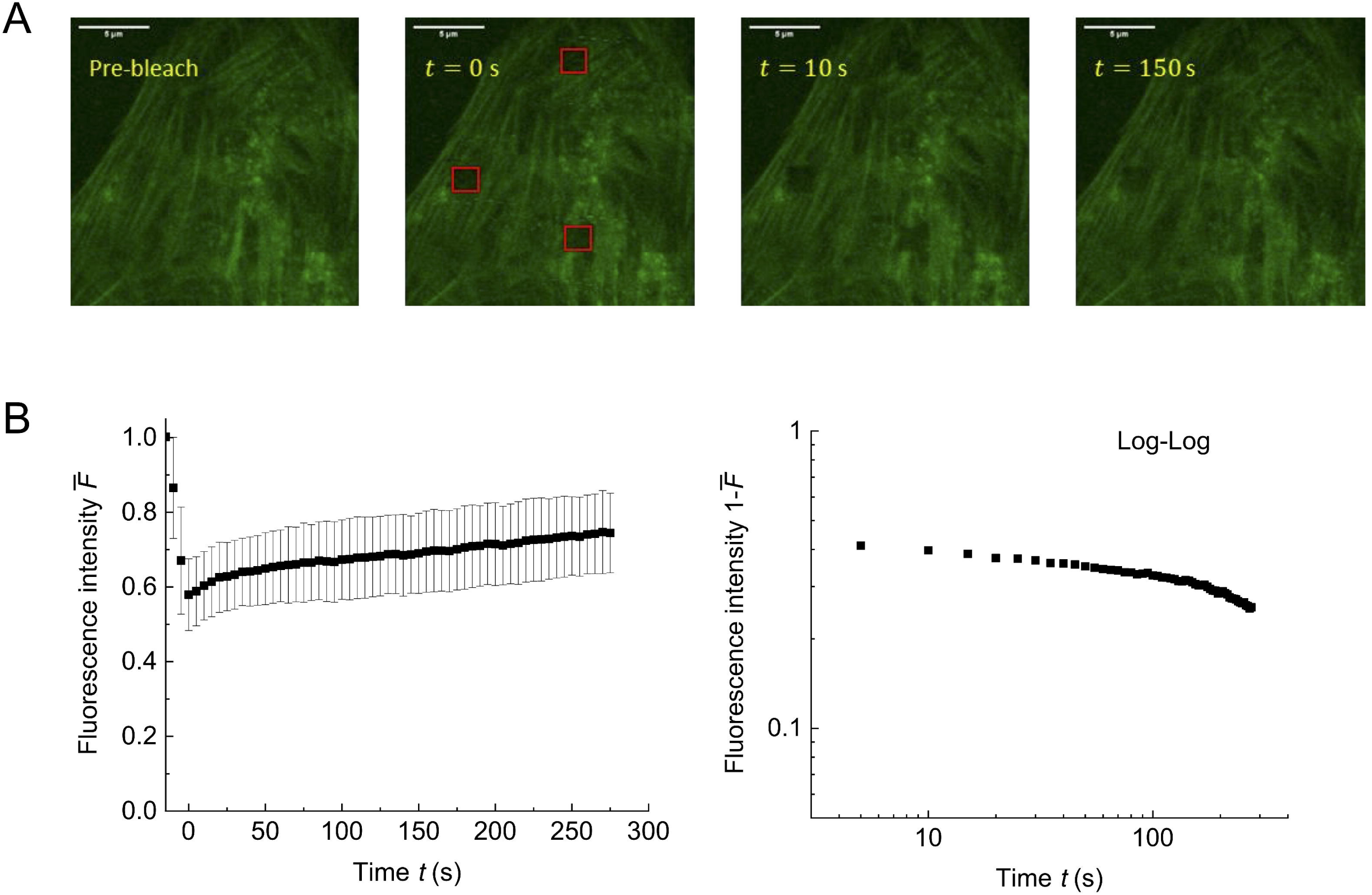
FRAP analysis suggests that F-actin in SFs works as an immobile scaffold for transgelin-2. (A) Confocal images of mClover2-beta-actin subjected to FRAP. Scale, 5 μm. (B) Time course of 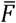 in the FRAP response of mClover2-beta-actin is shown by two ways – a linear scale plot (left, mean ± SD) and a log-log plot (right, mean) – showing that the recovery is obviously slower compared to that of transgelin-2 in Fig. 4.

### Distinction between the effective and pure diffusions enables the extraction of the reaction properties

To determine the effective diffusion coefficient *D*_eff_ based on Eq. (7), spatiotemporal recovery data *F*(*x,y,t*) up to the time that corresponds to the time constant *τ* was analyzed (Fig. 6A, Video S10). Compared to FL, *D*_eff_ was significantly increased for all the deletion mutants (Fig. 6B). The mutants without the CH domain – i.e., ΔCH, ΔCHΔCR, ΔABΔCH, and ΔABΔCHΔCR – all exhibited a markedly significant increase in *D*_eff_, likely because the greater reduction in their molecular weight compared to the deletion of the AB and CR domains might significantly elevate their diffusion rate, and/or the CH domain might critically contribute to the binding to actin compared to the AB domain. ΔCR resulted solely in a significant increase in *D*_eff_, which will be discussed below. In the analysis, photobleaching parameter and laser resolution were found to be ∼1 and ∼0.2 μm, respectively.

**Fig. 6.**
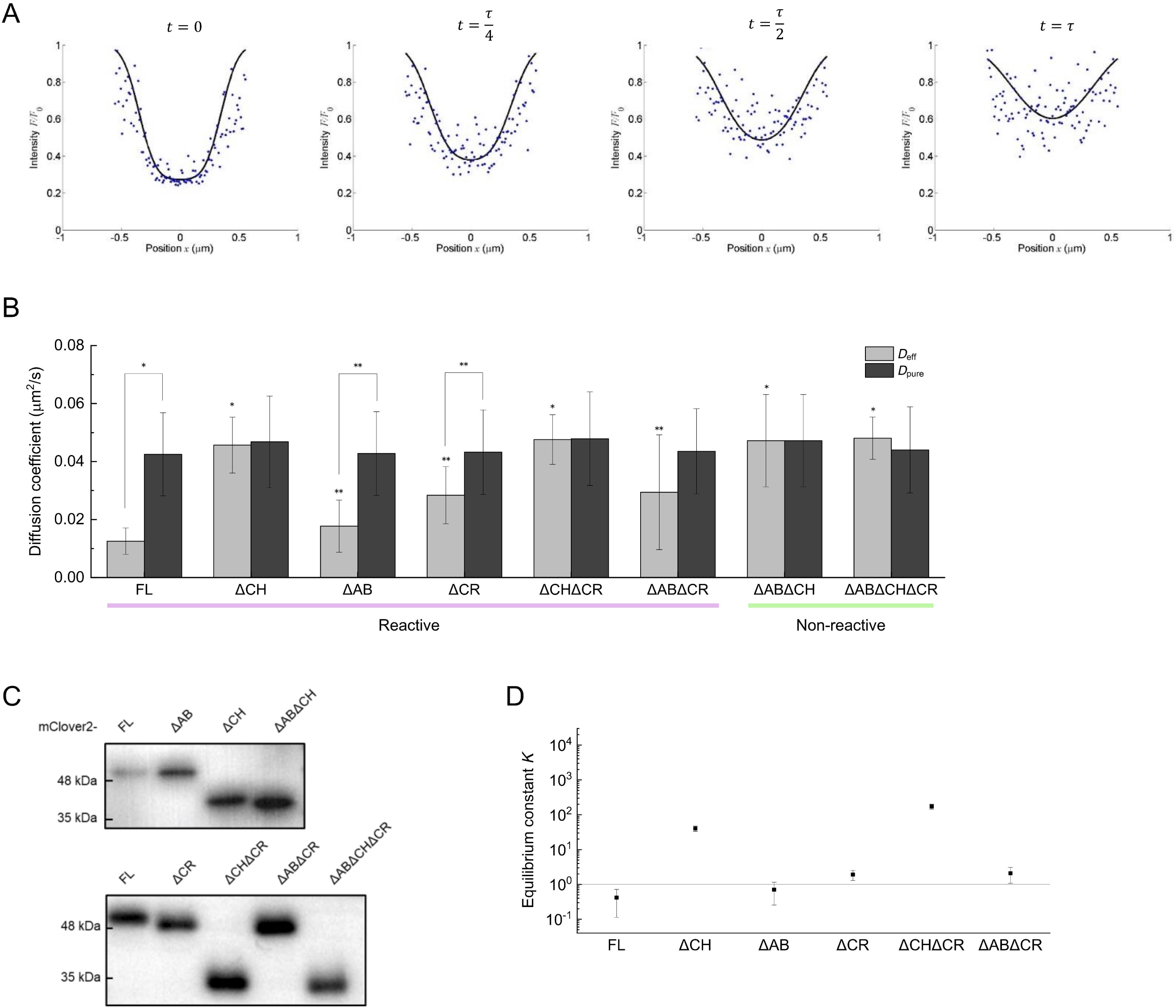
Individual domain-level determination of the reactive-diffusive properties. (A) Normalized fluorescence intensity *F*(*x, y* = 0, *t*)/*F*_0_ is plotted as blue dots over the position in the longitudinal *x* for specific time *t*, with the regression curves (black) determined by the maximum likelihood method. Here, representative data of FL are shown. *τ* represents the time constant, i.e., the inverse of the recovery rate *k* determined in Fig. 4. (B) *D*_eff_ (gray) and *D*_pure_ (black) for each mutant. Asterisks represent statistically significant differences compared to FL within the *D*_eff_ groups or between the specified pairs of *D*_eff_ and *D*_pure_ (*, *p* < 0.01; **, *p* < 0.05). (C) Immunoblots of transgelin-2 FL and its mutants with mClover2. (D) Equilibrium dissociation constant *K* for each mutant. Data are expressed as the mean ± coefficient of variation.

While *D*_eff_ of transgelin-2 was thus determined, the effective diffusion does not rigorously distinguish per se between pure diffusion and chemical interaction with other proteins. We then estimated the pure diffusion coefficient of transgelin-2 – which characterizes its inherent diffusive properties independent of the chemical interaction – from the specific *D*_eff_ determined in the non-reactive system, i.e., the mutants with no actin-interacting domain ΔABΔCH (Fig. 1B). To do this, we compensated for the reduction in the molecular weight due to the deletion of the domain(s) in the Stokes-Einstein equation to modify the pure diffusion coefficient in accordance with the reactive system. The molecular weight of each mutant with the fluorescent tag is known from the amino acid sequence (Fig. 3A), and indeed those values were consistent with the ones observed in the Western blots (Fig. 6C). In the non-reactive system, the effective diffusion coefficient is equivalent to the pure diffusion coefficient 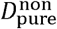, which was then converted to the pure diffusion coefficient in the reactive system 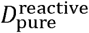 according to

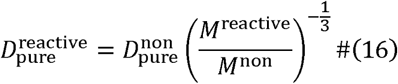

where *M*^non^ and *M*^reactive^ represent the molecular weight in the non-reactive and reactive systems, respectively. 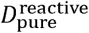 was thus obtained for each of the mutants as all the factors of the right side of Eq. (16) were now known (Fig. 6B). The acquired pure diffusion coefficients are relatively comparable in magnitude among the mutants because of the limited effect of the mass provided by one-third power of the ratio. Significant differences were found between the effective and pure diffusion coefficients in FL, ΔAB, and ΔCR, reflecting the contamination of the Brownian diffusion by the chemical interaction.

As partial validation, FCS was performed to assess the diffusion coefficient obtained by the above FRAP analysis although the measurable range of diffusion coefficients is typically ∼0.01–10 μm^2^/s for FRAP and ∼1–10^3^ for FCS (Matsuda and Nagai 2014; Lorén et al. 2015). The non-reactive mutant ΔABΔCH within SFs was estimated in the FRAP analysis to have a *D*_pure_ (= *D*_eff_) of ∼0.04 μm^2^/s (Fig. 6B), which is considerably below the effective range of FCS. Therefore, we instead focused on the same non-reactive mutant in the cytoplasm with looser actin meshwork, in which the molecules are supposed to diffuse faster compared to the inside of SFs composed of tighter actin bundles (Okamoto et al. 2020), and hence a larger *D*_pure_ closer to the range of FCS might be obtained. Our FRAP analysis thus determined *D*_pure_ of ΔABΔCH in the cytoplasm other than SFs to be 0.25 ± 0.14 μm^2^/s (Fig. S2). Meanwhile, FCS provided an estimate of 0.64 ± 0.16 μm^2^/s in the same conditions. *D*_pure_ of the representative molecule ΔABΔCH was thus shown to be approximately in the same order of magnitude regardless of the methods used, hence partially verifying the quantitative consistency with the other methodology and the plausibility of the other values determined based on our proposed FRAP framework.

Next, from the resulting sets of *D*_eff_ and *D*_pure_ (Fig. 6B), the equilibrium dissociation constant *K* = *k*_off_/*k*_on_ was obtained for both the reactive and non-reactive systems from Eq. (6) with the assumption that the chemical interaction takes place at equilibrium (Fig. 6D). A *K* value of the unity indicates a state that the association rate is equal to the dissociation rate, and thus the amount of the proteins is the same between the binding and unbinding states. FL with a *K* of <1 is thus suggested to be preferentially at the binding state. ΔCH and ΔCHΔCR with a *K* of ∼10–100 are considered to be predominantly at the unbinding state. ΔAB is only slightly higher in *K* compared to FL, suggesting that, while the AB domain is involved in the interaction with F-actin in SFs (Fig. 3B, C), the association via the AB domain is relatively unstable compared to that via the CH domain. ΔCR is also high in *K* compared to FL while this mutant still contains the actin-interacting domains of the AB and CH domains. It is likely that, while the CR domain of transgelin-2 is not directly involved in the association with F-actin, its deletion distorts the rest of the protein conformation to in turn reduce its affinity to F-actin, as previously indicated on the isoform transgelin-1 (Matsui et al., 2018). ΔABΔCR is consequently close to ΔCR in the average value of *K*.

## Discussion

To evaluate the turnover rate of proteins in FRAP experiments, it is often the case that only temporal change in the fluorescence intensity is analyzed, for example, using exponential functions like Eq. (1). On the other hand, to dissect the respective contributions of the two major factors – pure diffusion and chemical interaction with other molecules – to the overall “effective” diffusion, the spatial as well as temporal changes may be analyzed as we did here with Eq. (7). Importantly, one major challenge in the field of cell signaling research is to identify the role of specific domains of a protein in interacting with other molecules. To this end, the conventional approach in the FRAP analysis can be insufficient particularly in a case that the target protein has multiple domains to independently interact with the same types but different molecules. In this regard, transgelin-2 has been implicated in interactions with F-actin in cells putatively via two distinct domains (Castresana and Saraste 1995; Solway and Fredberg 1997; Na et al. 2015; Kim et al. 2018), making the intramolecular origin of the FRAP response uncertain, and was thus selected in the present study to test the validity of our new approach as well as to reveal the manner of the interactions.

We then developed a framework that incorporates comprehensive types of deletion mutants of transgelin-2 into spatiotemporal analyses of FRAP data. This approach requires some effort to construct the mutants, but instead clarifies the chemical equilibrium constants of intermolecular interactions at the individual domain level within the complex milieu of the cell. Pure diffusion properties, which are independent of the association/dissociation to the surrounding molecules and are thus intrinsic to the molecules, are also obtained in our analysis from the diffusive behavior of the non-reactive mutant. The described framework is universally applicable, in addition to transgelin-2 demonstrated here, to any protein bound to virtually immobile scaffolds like ventral SFs (Saito et al. 2020; Okamoto et al. 2020), thus providing a versatile approach to identifying the role of protein domains expressed within living cells.

Transgelin-2 has been implicated in various processes including tumorigenesis, cell migration/invasion, proliferation, differentiation, and embryo implantation (Yoshino et al. 2011; Yakabe et al. 2016; Jeon et al. 2018; Liang et al. 2019; Liu et al. 2019), but its mechanistic basis remains elusive. Transgelin-2 is known for its ability to polymerize G-actin and crosslink F-actin, which may eventually contribute – as many other ABPs that account for ∼25% of the total cellular protein do (Pollard 2016; Lappalainen 2016; Jo et al. 2018; Kim et al. 2018) – to maintaining the actin monomer pool available for the polymerization and promoting the elongation in a cell context-dependent manner. To quantitatively analyze these complicated cell processes in which a number of proteins are potentially involved, the knowledge obtained with simplified in vitro experiments where only a limited number of molecules are considered is not necessarily valid. In fact, the manner of the association of transgelin-2 to F-actin is distinct between in vitro and in cells. Specifically, among the known five types of CH domains, transgelin-2 has type-3 CH domain (Gimona et al. 2002) that is actually unable to bind F-actin in vitro (Gimona and Mital 1998; Fu et al. 2000; Goodman et al. 2003). Accordingly, the firmer association of the CH domain to F-actin over the AB domain within cells suggests that cellular transgelin-2 forms a complex with F-actin predominantly via a third protein bound to the CH domain. Given the presence of other actin-associated proteins containing type-3 CH-domain, such as calponin, vav, and IQGAP (Gimona et al. 2002; Yin et al. 2020), it is presumably difficult to accurately evaluate the intracellular physicochemical properties of at least these proteins by only conventional FRAP experiments. The in situ domain-level determination of the reaction properties independent of the pure diffusion nature will thus be useful, as presented here for transgelin-2, in revealing the real signaling mechanism of cellular proteins.

## Acknowledgments

TS is supported by Japan Society for the Promotion of Science (JSPS), Osaka, Japan.

## Supplementary materials

Fig. S1 Mobile fraction *A* in Eq. (1) for different mutants. Asterisks represent statistically significant differences compared to FL (*, p< 0.01).

Fig. S2 Quantified *D*_pure_ is comparable in magnitude between FCS and FRAP. (A) Normalized autocorrelation function *G*(*τ*)/*G*(0) of the fluorescence signal of the non-reactive ΔABΔCH in the cytoplasm (purple) as a function of the lag time *τ*. Background data are shown as a negative control (green). (B) Quantitative comparison of the *D*_pure_ of the non-reactive ΔABΔCH determined by the two distinct ways, FCS and FRAP. Boxes represent the 25th and 75th percentiles and the median. Open squares indicate the means. Whiskers extend from the box ends to the most remote points excluding the outliers defined as 1.5X the interquartile range. There is a significant difference between the two populations (*p* < 0.01), but they may be regarded comparable in order of magnitude, both with a *D*_pure_ value on the order of 0.1–1.0 μm^2^/s.

**Table S1.**
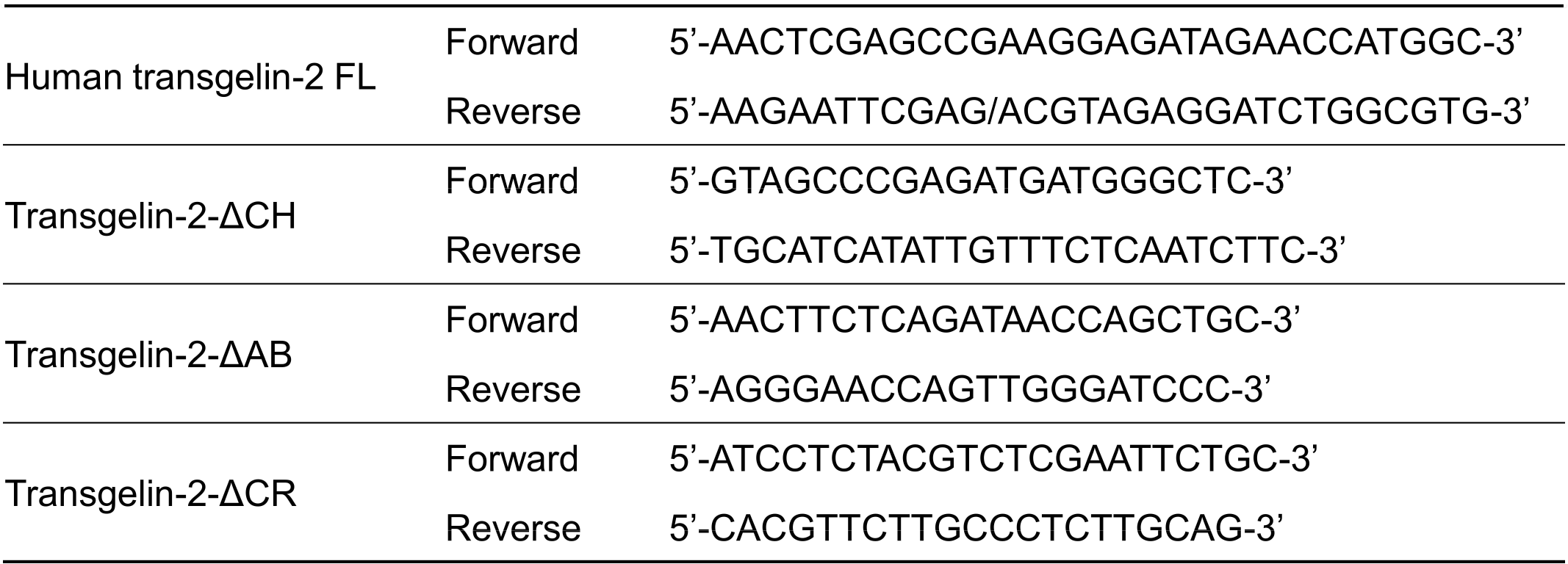
Primers for constructing transgelin-2 mutants.

**Table S2.** Parameters used for the conceptual analysis in Figs. 1A and 1B.

|  | Parameter | Value | Unit |
| --- | --- | --- | --- |
| Diffusion coefficient | $D_{\text{pure}}$ | 1 | $\mu\text{m}^2/\text{s}$ |
| Time step | $\Delta t$ | $10^{-2}$ | s |
| Number of particles | $N$ | $10^6$ | |

**Table S3.** Parameters used for the conceptual analysis in Figs. 1C, 1D, and 1E.

|  | Parameter | Value | Unit |
| --- | --- | --- | --- |
| Diffusion coefficient | $D_{\text{pure}}$ | $10^{-3}$ | $\mu\text{m}^2/\text{s}$ |
| Dissociation rate | $k_{\text{off}}$ | $5 \times 10^{-2}$ | $\text{s}^{-1}$ |
| Time step | $\Delta t$ | $10^{-3}$ | s |
| Increment | $\Delta x$ | $10^{-2}$ | $\mu\text{m}$ |
| Characteristic length | $L$ | 1 | $\mu\text{m}$ |
| Equilibrium concentration | $C_{\text{eq}}$ | 1 | M |

Video S1 Representative FRAP response of FL.

Video S2 Representative FRAP response of ΔCH.

Video S3 Representative FRAP response of ΔAB.

Video S4 Representative FRAP response of ΔCR.

Video S5 Representative FRAP response of ΔCHΔCR.

Video S6 Representative FRAP response of ΔABΔCR.

Video S7 Representative FRAP response of ΔABΔCH.

Video S8 Representative FRAP response of ΔABΔCHΔCR.

Video S9 Representative FRAP response of actin.

Video S10 Simulation of FRAP response (Fig. 6A).

